# Cell-type-specific execution of effector-triggered immunity

**DOI:** 10.1101/2025.06.28.662111

**Authors:** Himanshu Chhillar, Leonardo Jo, Amey Redkar, Kaisa Kajala, Jonathan DG Jones, Pingtao Ding

## Abstract

Effector-triggered immunity (ETI) is a central component of host defense, but whether all cell types execute ETI similarly remains unknown. We combined chemically imposed immune activation with single-cell transcriptomics to profile ETI responses across all leaf cell types in Arabidopsis. Despite uniform ETI perception, we find striking divergence between transcriptional outputs: a core set of defense genes is broadly induced, while distinct cell types activate specialized immune modules. We infer that downstream immune execution is shaped not only by immune receptor activation, but also by cell identity and its associated transcriptional regulatory context, including local transcription factor availability and chromatin accessibility. We further demonstrate that transcriptional regulators preferentially induced in epidermal cells are required to restrict invasion by non-adapted pathogens. Their absence permits pathogen entry into deeper tissues despite intact recognition, revealing a spatial division of immune functions. Our findings uncover a layered immune architecture in plants, challenges the assumption of uniform immune execution, and provides a framework for exploring cell-type-specific resistance logic in multicellular hosts.

## Introduction

In multicellular organisms lacking adaptive immunity, such as plants, immune responses are largely cell-autonomous: each cell must independently perceive and counter pathogenic threats in the absence of mobile immune cells^1^. While this principle has long shaped our understanding of plant defense, recent studies highlight the importance of intercellular communication in modulating local responses and coordinating systemic immunity^2–4^. Nevertheless, how immune execution at the resolution of individual cell types remains poorly understood. Specifically, it is unknown whether all plant cell types engage immune programs equivalently, or whether transcriptional responses are modulated by cell identity, developmental state, or physiological role.

Effector-triggered immunity (ETI), mediated by intracellular nucleotide-binding leucine-rich repeat (NB-LRR) receptors (NLRs), is a core component of plant innate immunity. Upon recognizing pathogen-derived effectors, ETI activates a robust transcriptional response that drives defense gene expression, cell wall fortification, immune metabolite production, and often culminates in a localized cell death^5^. While ETI is widely regarded as a canonical and system-wide immune program, most insights to date stem from bulk tissue measurements or natural infections, where immune activation is spatially heterogeneous and confounded by pathogen distribution and pattern-triggered immunity (PTI), mediated by cell-surface immune receptors. These limitations have obscured a fundamental question: Do all plant cell types execute ETI similarly when immune activation is synchronized?

Recent studies have begun to explore immune responses at single-cell resolution. For example, single-cell profiling of *Pseudomonas syringae*-infected Arabidopsis leaves revealed transcriptional shifts linked to pathogen proximity^6^, and fungal infection studies have identified cell-type-specific defense signatures under biotic stress^7^. Most recently, the concept of a primary immune responder (PRIMER) cell state was proposed, based on single-cell RNA-seq analysis of ETI-inducing bacterial infections during which host cells experience PTI, ETI and also defense suppression via pathogen effectors^8^. While these studies have uncovered valuable spatial features of plant immunity, they are inherently confounded by pathogen load, PTI co-activation, and the spatial unpredictability of natural infections.

To overcome these limitations, we established a synthetic system for uniform and pathogen-free ETI induction using estradiol-triggered NLR activation in Arabidopsis^9^. Combining this with single-cell RNA sequencing, we constructed a high-resolution atlas of ETI-responsive transcriptional states across major leaf cell types. Our analysis uncovered a layered immune architecture: despite of a shared core transcriptional response activated across all cell types, ETI responses were shaped by local gene regulatory networks in cell-type-specific transcriptional modules, demonstrating a complementary mechanism between both. This suggests that immune execution is not merely a product of perception but is also intrinsically constrained by cell identity and chromatin context.

To directly test whether cell-type-specific transcriptional modules are functionally required for immune execution, we focused on CAMODULIN-BINDING PROTEIN 60 G (CBP60g) and SYSTEMIC ACQUIRED RESISTANCE DEFICIENT 1 (SARD1), two closely related transcription factors known to act downstream of NLR signaling. Both have been established as central regulators of salicylic acid biosynthesis and systemic acquired resistance during ETI ^10–13^. Strikingly, our single-cell data revealed their preferential induction in pavement cells, the epidermal layer that interfaces directly with the external environment. This was particularly intriguing in the context of enabling a non-adapted *Albugo candida*, isolate to invade Arabidopsis through stomata that would normally be resisted. While guard cells form the stomatal pore, it is the surrounding pavement cells that must constrain early pathogen entry. We hypothesized that CBP60g and SARD1 may act as key mediators of spatial immune execution in this vulnerable layer. Supporting this, we found that *cbp60g sard1* mutants permitted haustoria formation by *A. candida* specifically in the epidermal and sub-epidermal cells, despite intact immunity in internal tissues. This provides direct evidence that cell-type-specific transcriptional activation, rather than receptor activation alone, is required to restrict nonhost pathogen ingress.

Together, our findings redefine our understanding of immune coordination in plants, not as a uniform switch, but as a mosaic of modular, spatially distributed programs embedded within tissues. By decoupling immune activation from pathogen-associated variables and resolving transcriptional responses at cellular scale, this work has broad implications for how multicellular hosts orchestrate defense and opens avenues for engineering more precise and spatially tuned immune activation in multicellular systems, including crops.

## Results

### ETI activation defines distinct immune-responsive cell clusters

To profile the transcriptional landscape of effector-triggered immunity (ETI) at single-cell resolution, we developed a synthetic ETI activation system in *Arabidopsis thaliana* (Arabidopsis thereafter) leaves using estradiol (E2)-inducible expression of *AvrRps4*, a bacterial effector recognized by the nucleotide-binding leucine-rich repeat (NB-LRR) receptor (NLR) pairs, RESISTANT TO RALSTONIA SOLANACEARUM 1 (RRS1)/RESISTANT TO PSEUDOMONAS SYRINGAE 4 (RPS4) and RRS1B/RPS4B, referred to as the SUPER ETI (SETI) line^9,14^. Leaves of the SETI plants were infiltrated with mock or E2 solution, and protoplasts were isolated 4 hours post-infiltration (hpi) for single-cell analysis (Figure 1A). Real-time quantitative PCR confirmed robust induction of one of the ETI early responsive marker genes, *ISOCHORISMATE SYNTHASE 1* (*ICS1*) (Figure 1B)^15,16^, validating effective immune activation.

**Figure 1.**
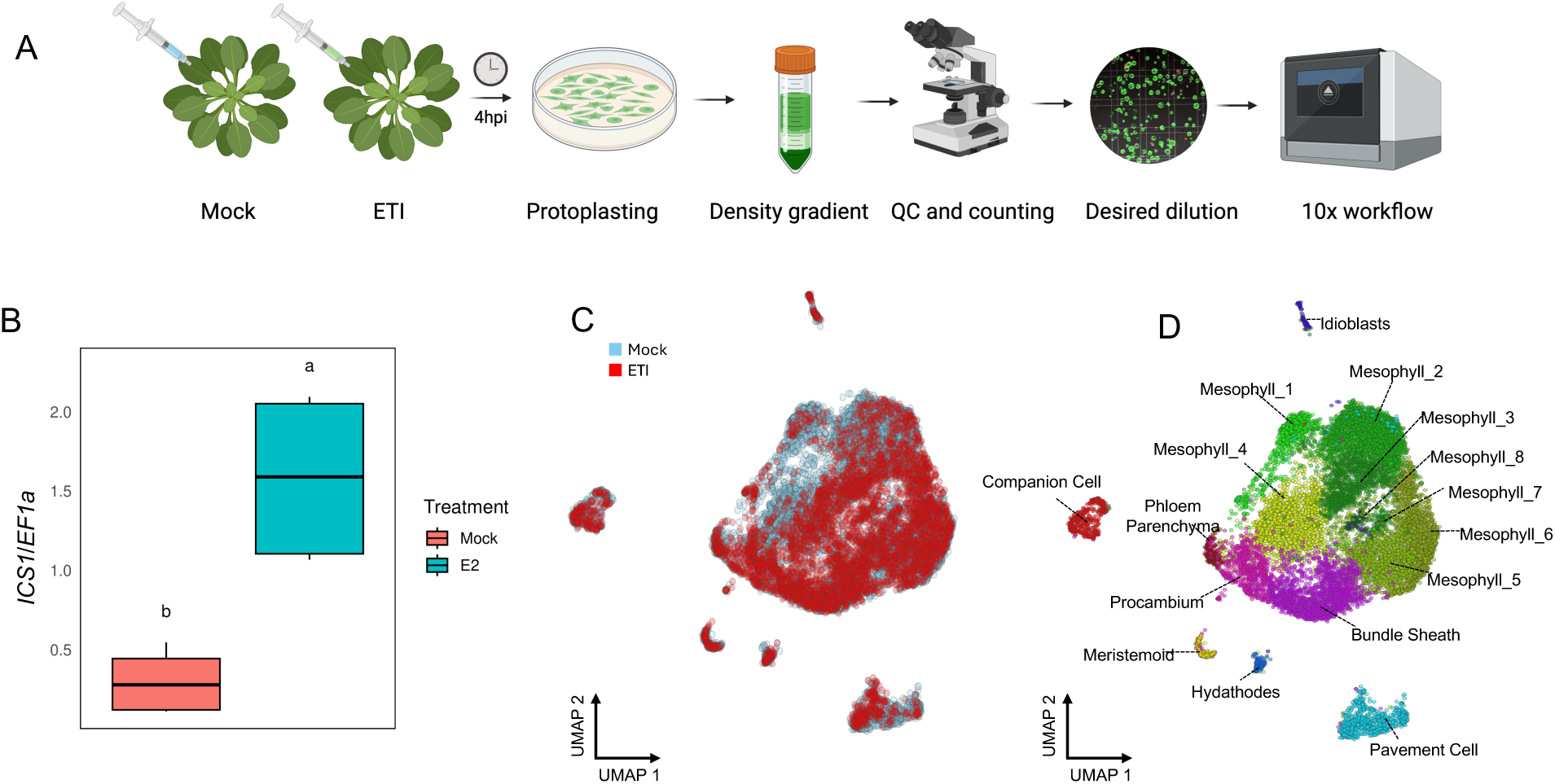
Single-cell transcriptomic landscape of immune activation in Arabidopsis leaf tissue. (A) Experimental workflow. Arabidopsis leaves were treated with mock or estradiol to induce effector-triggered immunity (ETI), followed by protoplast isolation at 4 hours post-infiltration (hpi) and single-cell RNA sequencing (scRNA-seq). (B) qPCR analysis of the immune marker gene *ISOCHORISMATE SYNTHASE 1* (*ICS1*). This confirms effective ETI induction in treated samples, validating immune activation prior to scRNA-seq library preparation. (C) UMAP projection of integrated single-cell transcriptomes from mock and ETI-treated samples, colored by condition. (D) UMAP showing unsupervised clustering of major Arabidopsis leaf cell types including pavement cells, mesophyll subtypes (Mesophyl_1 to Mesophyl_8), phloem parenchyma, companion cells, hydathodes, idioblasts, meristemoids, procambium, and bundle sheath cells.

We performed single-cell RNA sequencing (scRNA-seq) on protoplasts from mock and E2 treated samples using the 10X Genomics Chromium X platform and obtained approximately 300 million reads. After quality control, we recovered 7932 and 5739 high-quality cells in the mock- and E2-treated samples, respectively, with a median of 527 and 1134 genes and 776 and 2153 unique transcripts per cell, recovering approximately 70% of annotated protein-coding genes in the Arabidopsis genome. ETI activation led to notable changes in the relative abundance of specific cell types (Figure S2A), possibly reflecting global transcriptional reprogramming.

Unsupervised graph-based clustering identified 16 major cell clusters after integration of mock and E2 treated samples, visualized using uniform manifold approximation and projection (UMAP) (Figure 1C and 1D). Cells from both mock and E2 conditions were represented across clusters (Figure S1A and B). To assign cell identities, we leveraged a recently published single-cell atlas of Arabidopsis leaf tissue and the well-known canonical marker genes^17^. A violin plot of selected cell-type-specific genes (Figure S1C) confirmed robust classification of 16 clusters corresponding to nine distinct cell types, with a predominant mesophyll identity (clusters M1–M8) along with pavement cells (epidermal cells), bundle sheath, hydathode, phloem, and other known leaf cell types (Figure 1D).

Together, these data establish a comprehensive single-cell atlas of early ETI activation across Arabidopsis leaf tissue, in the absence of PTI, providing a foundation for dissecting immune responses at cellular resolution without confounding pathogen-derived heterogeneity or defense suppression.

### Uniform immune activation elicits heterogeneous transcriptional responses

To dissect the spatial pattern of immune execution, we examined the expression of canonical immune and susceptibility genes across annotated cell types. Core components of the Toll/Interleukin-1 receptor/Resistance protein (TIR) NLR immune signaling cascades including *ENHANCED DISEASE SUSCEPTIBILITY 1* (*EDS1*), *PHYTOALEXIN DEFICIENT 4* (*PAD4*) and *SENESCENCE ASSOCIATED GENE 101* (*SAG101*) were broadly expressed, along with downstream helper NLR encoding genes, *ACTIVATED DISEASE RESISTANCE 1* (*ADR1*), *ADR1-L1, ADR1-L2*, *N REQUIREMENT GENE 1.1* (*NRG1.1*) and *NRG1.2* (Figure 2A)^18^, with *ADR1* and *NRG1.2* exhibiting relatively sparse expression. This widespread expression suggests that the EDS1-dependent module is transcriptionally accessible across the entire leaf tissue, equipping most cell types to mount ETI responses upon intracellular effector recognition.

**Figure 2.**
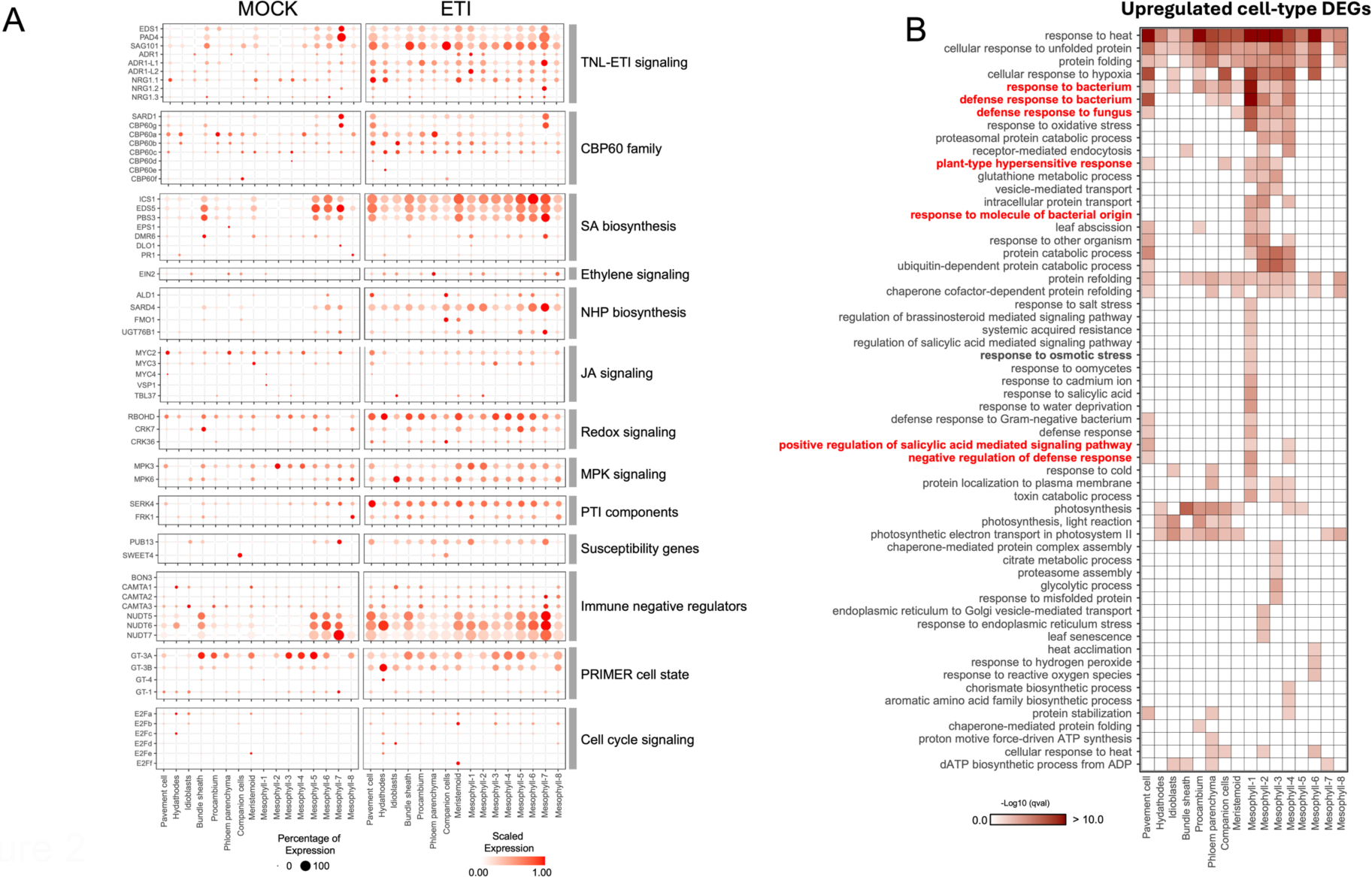
Cell-type-specific transcriptional responses to ETI in Arabidopsis leaf tissue. (A) Bubble plot showing expression of representative immune-related genes across annotated cell types under mock and E2 conditions. Categories include TNL-ETI signaling components, CBP60 family members, salicylic acid (SA), jasmonic acid (JA), ethylene, and NHP signaling, redox and MAP kinase pathways, PTI components, susceptibility factors, and immune repressors. (B) Gene Ontology (GO) enrichment of upregulated DEGs in ETI-treated cells. Functional categories include defense responses, hormone signaling, oxidative stress, protein folding, photosynthesis, and vesicle-mediated transport. Enrichment significance is shown as –log10(q-value), illustrating spatially resolved immune reprogramming during ETI.

Consistently, the phytohormone salicylic acid (SA) biosynthesis pathway^19^, downstream of EDS1 signaling, was also globally activated. Genes including *ICS1*, *EDS5* and *AVRPPHB SUSCEPTIBLE 3* (*PBS3*) were robustly expressed across nearly all cell clusters (Figures 2A, S2B and S2C)^19^, in line with broad SA pathway activation previously observed in ETI^16,20^. The widespread induction of *ETHYLENE INSENSITIVE 2* (*EIN2*) further supports the notion of a tissue-wide immune readiness (Figure 2A). Notably, negative immune regulators such as Nudix hydrolase encoding genes*, NUDTs*, and *CALMODULIN-BINDING TRANSCRIPTION ACTIVATORs* (*CAMTAs*) were also broadly induced (Figure 2A), indicating that immune repression mechanisms are similarly embedded across the tissue.

Moreover, key signaling components that mediate the mutual potentiation between PTI and ETI, including shared protein kinases and transcriptional regulators^14^, were also ubiquitously expressed (Figure S3A). This suggests that all cells in Arabidopsis leaves harbor an immunity-supporting infrastructure that is transcriptionally primed to interpret and integrate diverse immune cues.

In contrast to these broadly distributed immune modules, we identified marked spatial restriction in the expression of N-hydroxy pipecolic acid (NHP) biosynthesis genes. Specifically, *AGD2-LIKE DEFENSE RESPONSE PROTEIN 1* (*ALD1*) and *FLAVIN-CONTAINING MONOOXYGENASE 1* (*FMO1*), which encodes enzymes catalyzing the first and last steps in NHP production^21–23^, were almost exclusively expressed in companion cells (Figures 2A, and S2D and S2E), consistent with recent findings^8^. Surprisingly, *SYSTEMIC ACQUIRED RESISTENCE DEFICIENT 4* (*SARD4*), which encodes the oxidoreductase catalyzing the intermediate step of NHP biosynthesis^21,24^, exhibited widespread expression (Figure S2F), suggesting a division of labor wherein companion cells complete NHP biosynthesis to support systemic signal propagation. These findings indicate a spatial configuration of NHP biosynthesis and reinforce the role of companion cells as specialized immune hubs for systemic acquired resistance (SAR)-related metabolites.

We also observed cell-type-restricted expression for genes in the jasmonic acid (JA) signaling pathway and members of the E2F transcription factor family (Figure 2A), previously implicated in cell cycle control and immune regulation^15,25,26^. These results suggest that hormone signaling, and proliferation-linked regulators may modulate ETI output in a spatially restricted and context-dependent manner. For instance, loss-of-function of *E2Fa/b/c* compromises ETI responses mediated by both TIR and coiled-coil (CC) NLRs^26^.

Importantly, because our system triggers ETI uniformly and in a pathogen-free manner^9^, it offers a unique opportunity to examine whether immune-enhanced cell states, such as primary immune responder (PRIMER) and bystander cell states^8^, can arise independently of pathogen spatial distribution. We observed broad induction of the trihelix transcription factor encoding gene, *GT-MOTIF BINDING FACTOR-3A* (*GT-3A*)^27^, a canonical marker of the PRIMER cell state^8^, across all ETI-activated cell types (Figure 2A), consistent with its role as a general ETI-responsive gene. This confirms that *GT-3A* expression reflects the proximity to pathogens that can induce robust cell-autonomous ETI. In contrast, its close homologues *GT-3B* and *GT-4* were preferentially induced in hydathodes (Figure 2A), indicating cell-type-specific modulation of the transcriptional responses mediated by putatively redundant components. These findings refine the PRIMER cell state concept by suggesting that its transcriptional hallmarks can be broadly accessible to almost all plant cells but differentially tuned by cell identity. Rather than being exclusively driven by pathogen contact, PRIMER-like transcriptional cell states may emerge from intrinsic regulatory programs that govern immune execution across tissues.

Taken together, these results reveal that while the capacity to initiate immune signaling is broadly encoded in all leaf cells, the execution of immune programs is highly modular and spatially patterned. Even under synchronized and pathogen-free ETI activation, immune outputs such as NHP biosynthesis and PRIMER-like transcriptional states remain confined to specific cell types. This indicates that plant tissues harbor intrinsic immune compartmentalization, likely reflecting functional specializations that balance defense intensity, metabolic cost, and coordination of systemic immunity. These insights offer a new framework to explore how spatial immune signaling is uncoded in development and potentially exploited during natural infections.

### Shared and cluster-specific gene programs structure immune execution

To map the cellular architecture of immune execution, we performed differential gene expression (DEG) analysis across all annotated cell clusters (Figure S3B), followed by gene ontology (GO) term enrichment. As anticipated, defense-related GO terms were predominantly enriched in epidermal and mesophyll cells (Figure 2B, Table S2), consistent with prior bulk RNA-seq findings^16^. Interestingly, GO terms related to photosynthesis, especially photosystem I components, were significantly enriched among downregulated genes in several mesophyll clusters (Figure S4A), indicating that ETI alone can suppress core metabolic programs in addition to activating defense.

To evaluate how cell-resolved responses relate to canonical ETI programs, we compared upregulated DEGs in each cluster with a previously defined list of EDS1-dependent immune-responsive genes (Table S3)^16^. Most clusters showed significant overlap, with pavement cells and mesophyll cells exhibiting the highest concordance (Figure 3A and S4B). This supports the presence of a broadly conserved core immune program across major cell types.

**Figure 3.**
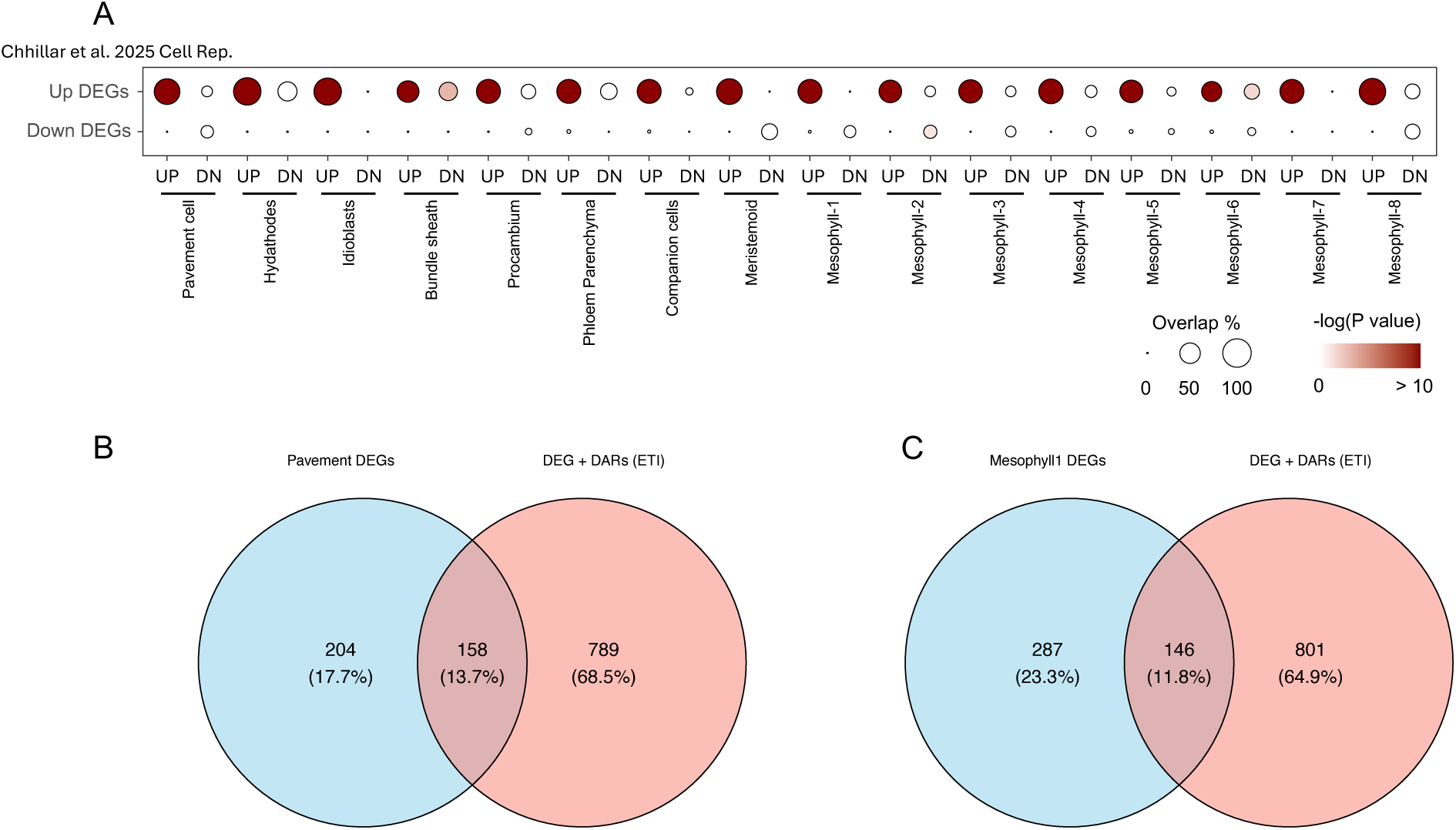
Integration of bulk RNA-seq and chromatin accessibility data with cell-type-specific ETI responses. (A) Hypergeometric test showing the significance of overlap between bulk ETI-responsive genes (Chhillar et al. 2025, *Cell Rep.*) and DEGs identified in individual cell clusters from single cell RNA-seq. (B–C) Venn diagrams showing overlap between DEGs and genes associated with differentially accessible regions (DARs) during ETI (from Ding et al. 2021, *JXB*) in pavement cells (B) and mesophyll-1 cells (C). A majority of cell-type-specific DEGs are also associated with DARs, suggesting coordinated transcriptional and chromatin-level regulation. Percentages reflect the proportion of DEGs overlapping with DARs in each cell type.

However, beyond this shared module, many cell clusters displayed distinct expression of immune-activation indicators or regulators, such as *CALMODULIN-BINDING PROTEIN 60 G* (*CBP60g*), *SARD1*, *ALD1*, *FMO1*, *SWEET4* and *E2F* family transcription factors etc., which may mediate cell-autonomous immune amplification, niche-specific defense execution, or immune signal propagation contributing to pathogen restriction and distal immune priming signal propagation.

To delineate these features systematically, we classified DEGs as shared (included in more than 2 different clusters; Table S7) or cluster-specific (uniquely induced; Table S8). Many cluster-specific genes still overlapped with bulk-identified ETI targets, suggesting these are not artifacts of cellular noise but reflect specialized, cell type-enriched immune indicators and putative regulators. Additionally, these genes overlapped with known EDS1-dependent ETI-responsive genes from bulk RNA-seq, supporting their role as specialized, cell type-enriched immune markers or execution components. Together, these results define a layered transcriptional framework for ETI in Arabidopsis: a core set of conserved immune genes is broadly deployed, while modular, cell-specific programs likely tailor immune outputs in a spatially precise manner. This modular immune architecture likely enables plants to coordinate tissue-scale defense while optimizing energy use and in parallel restrict pathogen invasion.

### Gene regulatory networks orchestrate cell-type-specific immune outputs

To elucidate how a uniform ETI activation translates into spatially heterogeneous immune execution, we examined the underlying regulatory mechanisms at the transcriptional level. We intersected upregulated DEGs from pavement and mesophyll 1 clusters with genes previously shown to be both ETI-induced (bulk RNA-seq) and exhibit ETI-enhanced chromatin accessibility (Table S4)^20^. This revealed a core set of chromatin-licensed immune genes (Figures 3B and 3C; Table S5 and S6), likely under transcriptional control during early ETI, and broadly conserved across both pavement and mesophyll 1 cell types.

To further dissect upstream regulatory control, we mapped cell-type specific DEGs to the Arabidopsis transcription factor (TF) *cistrome* and *epicistrome* database^28^. The *cistrome* defines the genome-wide landscape of transcription factor binding, while the *epicistrome* extends this by incorporating epigenetic features, such as DNA methylation, that modulate TF-DNA interactions, thereby shaping functional access to regulatory DNA. Together, they provide a comprehensive view of how gene regulatory networks are both encoded and epigenetically tuned^28^. WRKY, NAC, ERFs, MYB and bZIP TFs were prominently enriched in our analysis, consistent with established roles in plant immunity and suggesting that shared TF modules are deployed in distinct configurations across cell types to shape local defense outputs (Figure 4A)^29^.

**Figure 4.**
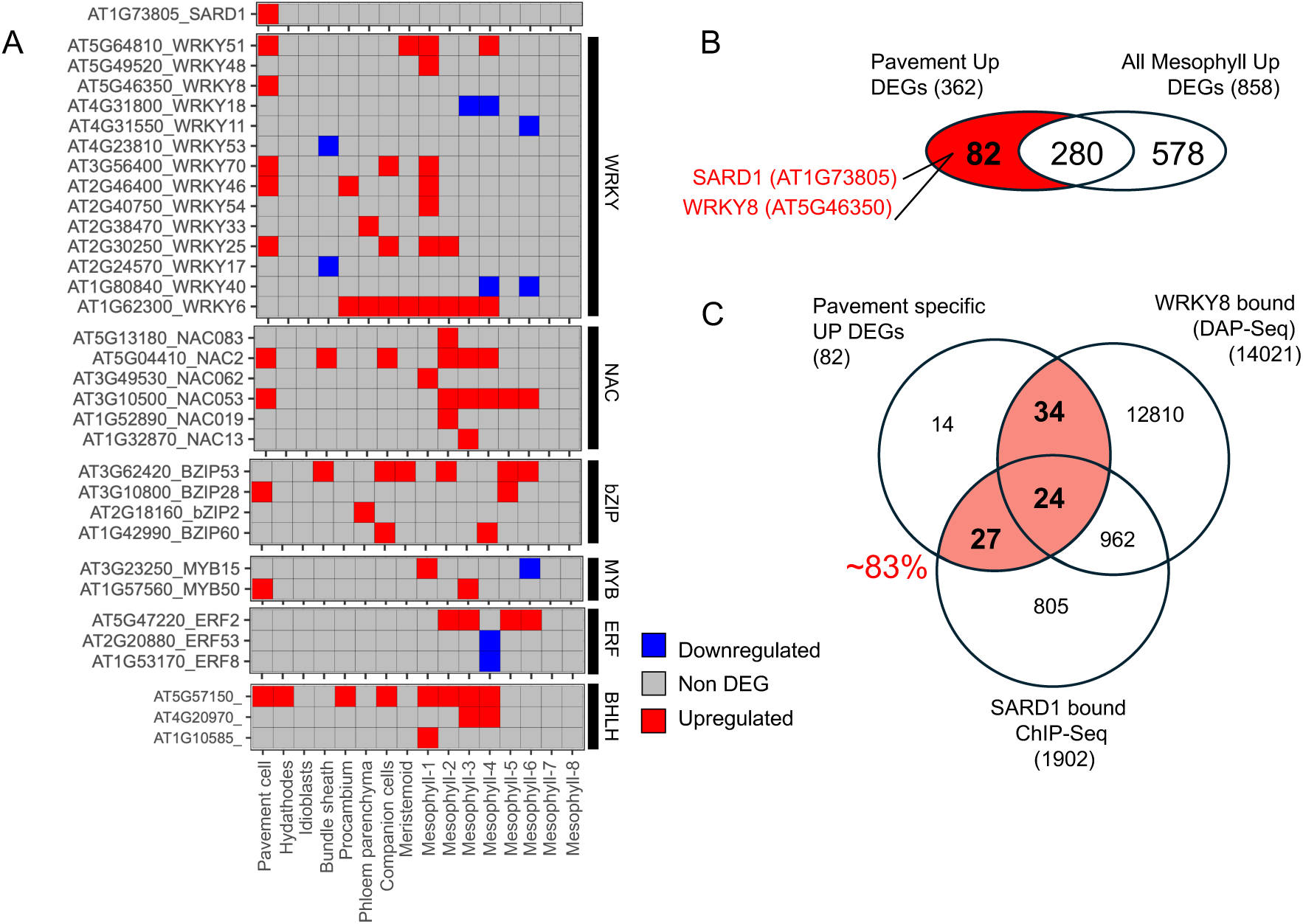
Cell-type-specific transcription factor activity underlies spatial immune regulation. (A) Heatmap showing differential expression patterns of selected transcription factors (TFs) across annotated Arabidopsis leaf cell types. TFs are grouped by family (WRKY, NAC, bZIP, MYB, ERF, and bHLH). Red and blue indicate up- and downregulated expression, respectively, under ETI. Each column represents an individual cell type. (B) Venn diagram comparing upregulated DEGs in pavement versus mesophyll cells. A subset of 82 genes is uniquely upregulated in pavement cells, including *SARD1* (*AT1g73805*) and *WRKY8* (*AT5g46350*). (C) Overlap between pavement-specific upregulated DEGs and genomic targets of WRKY8 (DAP-seq) and SARD1 (ChIP-seq). Approximately 83% of pavement-specific DEGs are putative direct targets of WRKY8 and/or SARD1, indicating coordinated transcriptional control of epidermal immune programs.

Notably, the enriched TFs exhibited varied responses across cell types, reflecting diverse regulatory roles in immune activation. Pavement and mesophyll cells showed the highest number of DEGs and the strongest GO enrichment for immune-related processes. We therefore compared these two cell types to investigate transcriptional regulators underlying spatial specificity.

Strikingly, *WRKY8* and *SYSTEMIC ACQUIRED RESISTANCE DEFICIENT 1* (*SARD1*) were both exclusively expressed in the epidermis (Figure 4A and 4B). Mapping to the *cistrome* and *epicitrome* database, WRKY8 was enriched at the promoters of a large fraction of pavement-specific ETI-induced DEGs (Figure 4C), consistent with its proposed role as an epidermal immune amplifier (Figure 4B)^8^. SARD1, a master regulator known to act redundantly with CBP60g to regulate plant immunity^10,11^, was similarly epidermis-specific and targeted more than 80% of upregulated DEGs in this cell type (Figure 4C). Together, these TFs accounted for the majority of regulatory activity in the epidermal immune network, underscoring their importance in orchestrating cell type–specific defense programs. Similar modular patterns were observed for other TFs with respect to other cell-types (Figure 4A), reinforcing idea that different TF families contribute to spatially specialized transcriptional circuits.

Collectively, these findings support a model in which ETI activation is interpreted through a preconfigured, cell-intrinsic transcriptional framework, governed by chromatin accessibility and transcription factor availability. This spatially modular regulatory architecture allows for the diversification of immune outputs despite a common upstream signal, enabling plants to finely tune defense intensity, coordinate intercellular signaling, and maintain metabolic balance under immune stress or during pathogen attack.

### Epidermis-specific immune regulators restrict non-host pathogen entry at the leaf surface

While many immune-related genes were broadly expressed across all leaf cell types upon ETI induction, certain transcriptional regulator exhibited strikingly cell-type-specific induction patterns, suggesting functional compartmentalization in immune execution. Among these, *CBP60g* and *SARD1*, two well-established, partially redundant immune activators^10,11,13^, were preferentially more upregulated in pavement cells (Figure 2A). SARD1 is a DEG specific to only pavement cells (Figure 4A), indicating a potential role in epidermis-localized defense mediated by ETI.

To assess whether this spatial enrichment reflects a functional requirement for immunity, we challenged *cbp60g sard1* double mutants and wild-type (WT) Col-0 with *Albugo candida* isolate AcEm2, a non-adapted oomycete pathogen that exhibits an incompatible interaction with Col-0 due to the NLR WHITE RUST RESISTANCE 4 (WRR4)-mediated recognition^30–32^. In contrast to WT plants, *cbp60g sard1* mutants permitted haustorial formation specifically in the epidermal and sub-epidermal cell layers, by not in deeper mesophyll tissues (Figure 5A and 5B). This indicates that while CBP60g and SARD1 are not globally required for ETI, they play a cell-type-specific role in restricting early pathogen entry at the leaf surface.

**Figure 5.**
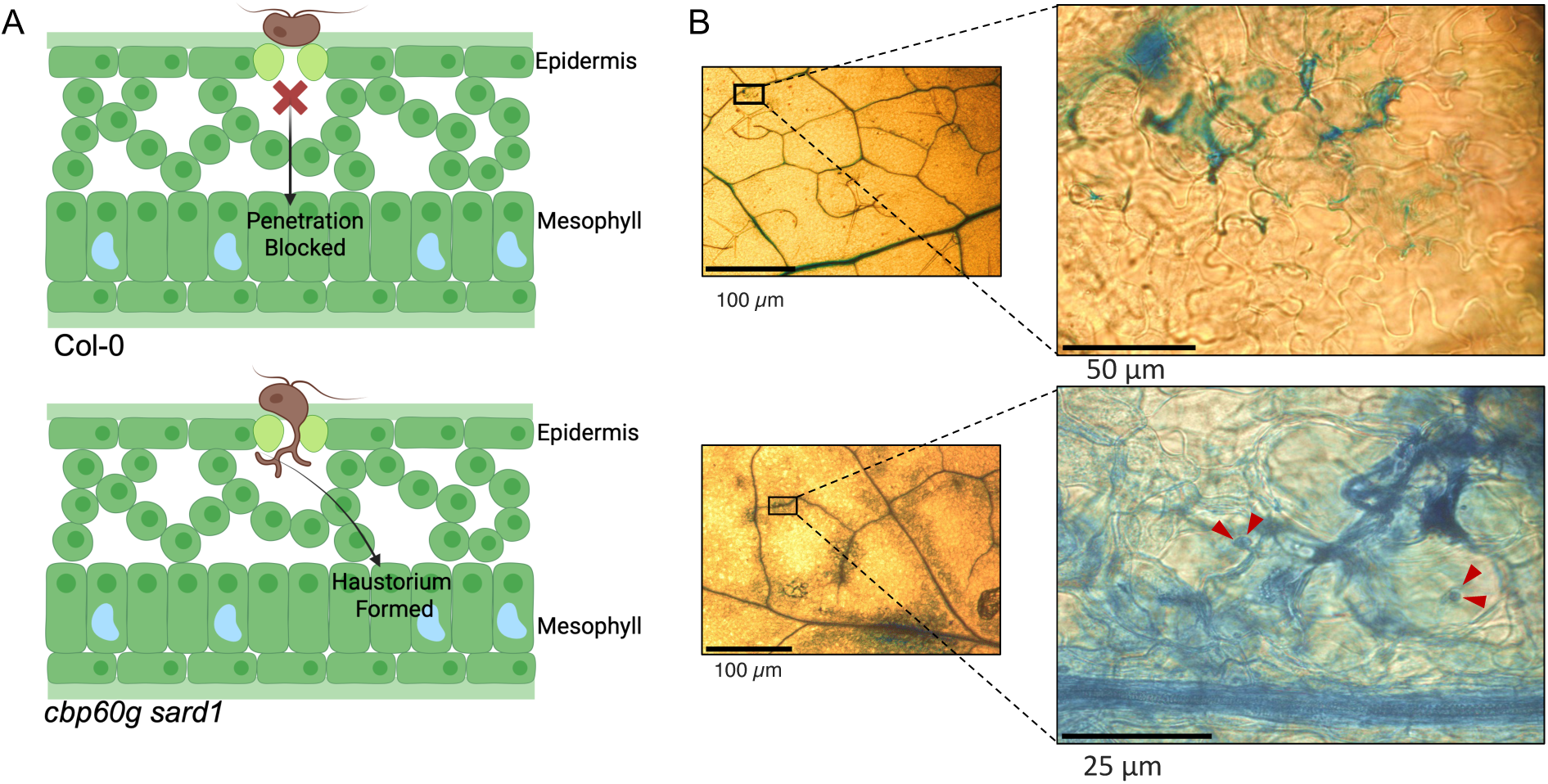
CBP60g and SARD1 mediate epidermis-specific immunity that *Albugo candida* **entry.** (A) Schematic representation of infection outcomes in *Arabidopsis thaliana* Col-0 and the *cbp60g sard1* double mutant following inoculation with *Albugo candida* isolate AcEm2. In Col-0, pathogen entry is blocked at the epidermal layer, whereas *cbp60g sard1* mutants permit haustorium formation in underlying mesophyll cells, indicating compromised epidermal defense. (B) Microscopic images of trypan blue-stained leaves infected with *A. candida* AcEm2. In Col-0, infection is restricted to the epidermis with no visible haustoria. In contrast, *cbp60g sard1* leaves show extensive hyphal growth and haustoria (red arrowheads) penetrating into mesophyll tissues, consistent with the model shown in (A).

Despite their loss, ETI signaling in mesophyll cells appears intact, likely due to broad expression of core components such as EDS1, PAD4, and other transcriptional regulators like WRKY33 or E2Fs that may independently sustain immune competence in internal tissues. This reinforces the concept that cell-type-specific TF modules mediate immune execution in distinct spatial zones, with CBP60g and SARD1 acting as epidermis-enriched executors to enforce localized resistance against non-adapted pathogens. While we did not perform pathogen-infected single-cell profiling, our data provide a transcriptional reference framework for immune potential across cell types and highlight testable hypotheses for future spatial challenge assays.

Furthermore, testing additional pathogens with alternative invasion routes, such as *Blumeria graminis*, which penetrates directly through the epidermis^33^, may clarify whether CBP60g- and SARD1-dependent immunity represents a generalizable outer-layer defense strategy. Such studies will further refine our understanding of how immune competence and execution are partitioned across tissue layers to collectively enforce nonhost resistance.

## Discussion

Our study presents a single-cell transcriptional atlas of ETI in Arabidopsis leaves, enabled by the SETI system^9^, which uncouples immune activation by pathogen-derived cues (*e.g.* Mechanical perturbations, cell-wall integrity and apoplastic chemical matrix alternations, and cell-surface receptor-mediated signaling upon elicitor recognition, etc.) from the effects of pathogen effectors. This approach reveals how intracellular ETI is broadly perceived yet executed in a spatially heterogeneous manner across different cell types, offering new insights into the modular logic of plant immunity.

ETI activation via TIR-NLRs induces a widespread transcriptional competence across leaf cell types. Genes encoding core components such as EDS1, PAD4, SAG101 and enzymes involved in salicylic acid (SA) biosynthesis are broadly expressed, indicating that immune signal transduction is not restricted to a specialized subset of cells within the leaf, but is accessible to diverse cell types. Nonetheless, specific immune outputs, including the NHP and JA pathways, are spatially restricted, suggesting compartmentalization of defense metabolism. These results imply that metabolic branches of ETI may follow cell-type-specific coordination, raising the need for future integration of single-cell metabolomics to map the immune metabolome with cellular precision.

At the gene regulatory level, our data support a layered and modular tissue architecture of immune control. While certain transcriptional modules are shared across cell types, others are distinctly partitioned. WRKY25 and WRKY70 exhibit broad deployment, whereas WRKY8 appears to function almost exclusively in pavement cells, regulating the majority of immune-induced genes in that cellular layer. Such spatial exclusivity is consistent with proposed role of WRKY8 in the PRIMER cell state. However, our results challenge the notion of the PRIMER state as a rare phenomenon. Instead, we propose that it represents a manifestation of differential ETI execution programs across cell types, embedded in their intrinsic components such as chromatin landscapes and transcription factor availability. The observed spatial heterogeneity reflects not a stochastic occurrence, but a structured immune regulatory mechanism encoded in tissue architecture. Other TFs, including WRKY17, WRKY33, E2Fs, and trihelix factors, further reinforce the diversity of transcriptional programs orchestrating immune output across spatial domains. This emphasizes that transcriptional immune competence is not synonymous with uniform immune execution. Despite shared upstream signaling through TIR-NLRs and EDS1, each cell type interprets immune signals through its own transcriptional lens, shaped by chromatin accessibility, TF availability, and cellular identity. This modular state likely allows plants to fine-tune defense intensity, optimize resource allocation, and coordinate systemic signaling in a spatially controlled manner.

Conventionally, nonhost resistance (NHR) represents a durable and broad-spectrum form of plant immunity, in which an entire plant species is resistant to all strains of a non-adapted pathogen^34^. While traditionally attributed to preformed barriers and pattern-triggered immunity (PTI), recent studies suggest that intracellular nucleotide-binding leucine-rich repeat (NLR) receptors can also contribute to nonhost restriction, particularly through effector recognition and ETI-like responses in specific pathosystems^35^. For instance, Arabidopsis NLR WRR4A mediates resistance to multiple isolates of *Brassica*-infecting isolates of *Albugo candida* ^36^, by recognizing lineage-specific effectors and triggering localized immune activation^30–32^. Our results extend this model by demonstrating that spatially restricted transcriptional regulators such as CBP60g and SARD1 are essential for enforcing cell-type-specific resistance against non-adapted *Albugo* isolates, preventing pathogen ingress at the epidermal layer. In *cbp60g sard1* double mutants, *A. candida* haustoria are formed in cell types at the outer layer of the leaf tissue, despite intact immunity in underlying tissues. This provides direct evidence that immune competence alone is insufficient without proper transcriptional execution at vulnerable interfaces. It also underscores a broader principle: spatially localized immune regulators enforce layered defense barriers that guard vulnerable tissue interfaces, a robust framework that could likely be conserved across diverse plant-pathogen interactions. Building on this, future studies could extend similar analyses, combined with spatial omics approaches, to identify additional spatially enriched immune components and evaluate their cell type–specific functional relevance.

The hydathode, a specialized structure at the leaf margin that connects to the vasculature, exemplifies this principle. Unlike the epidermis, which confronts external pathogens, hydathodes represent potential internal entry points. Prior work has shown that hydathode immunity is mediated by EDS1-PAD4-ADR1 signaling^37,38^, and our findings extend this by identifying GT-family transcription factors and GT-3A paralogs, GT-3B and GT-4, as selectively induced in hydathodes during ETI. While GT-3A is expressed in almost all cell types during ETI activation, the restricted expression pattern of GT-3B and GT-4 suggests that these TFs constitute a hydathode immune regulatory module. This underscores how even uniformly activated ETI is interpreted through distinct local transcriptional regulation, shaped by structural vulnerability and functional necessity.

Our findings deliver several conceptual advances. First, we provide direct transcriptional evidence that innate immune competence is broadly encoded in all major cell types of the Arabidopsis leaf, supporting a model of fully distributed, cell-autonomous immunity. Second, we show that even under uniform activation of intracellular immunity, cell identity exerts a major constraint on the spectrum and strength of transcriptional output. This introduces the concept of cell-type-specific immune execution, highlighting new questions about how chromatin features, TF availability, and metabolic state shape these responses. Our dataset offers a valuable resource for future mechanistic studies into these layers of regulation. Third, our functional validation of the epidermis-enriched transcriptional regulators CBP60g and SARD1 establishes that NHR mediated by NLR-triggered ETI is not uniform across tissues but depends on precise execution in specific cell types. In *cbp60g sard1* double mutants, haustoria of resisted races of oomycete *A. candida* formed specifically in the epidermal and sub-epidermal cells, despite intact immunity in inner layers. This demonstrates that immune competence alone is insufficient without proper transcriptional execution at vulnerable tissue interfaces. These findings provide, to our knowledge, the first direct evidence that NLR-driven NHR displays cell-type specificity, with functional consequences for pathogen entry.

Altogether, our results define a spatially modular model of ETI, where uniformly initiated immune signaling is filtered through localized transcriptional programs. This allows for precise deployment of immunity, which is stronger in vulnerable cell types in Arabidopsis leaf tissue like the epidermis and hydathodes than elsewhere. These insights raise compelling research questions: What molecular features prime certain cells for heightened immune execution? Can spatial chromatin states predict immunocompetence? How do localized defenses coordinate with systemic signaling? Is infection spatially random, or shaped by host–pathogen combinations at the single-cell level? How do highly responsive cells compensate for neighboring cells with limited immune potential to maintain tissue-wide resilience?

In summary, this work provides a foundational framework for understanding cell-type-resolved immunity in plants. By demonstrating how immune signaling is broadly competent yet differentially executed, we reveal the principles of spatial immune execution. These insights offer a roadmap for engineering durable, precision immune responses in crops, tailored to the distinct roles and vulnerabilities of each cell type.

## Methods

### Plant material and growth conditions

*Arabidopsis thaliana* Col-0 ecotype was used as the wild-type background. The β-estradiol (E2)– inducible SUPER-ETI line (SETI_WT) and the *cbp60g sard1* double mutant used in this study have been previously described^9,10^. Plants were grown at 21°C under long-day conditions (16 h light, 8 h dark), and at 50% humidity.

#### Protoplast isolation

Four-week-old leaves from SETI_WT plants were infiltrated with 50 μM estradiol (E2), and protoplasts were isolated 4 hours post-infiltration (hpi) using the Tape-Arabidopsis Sandwich method^39^. Peeled leaf tissue was incubated in enzyme digestion solution containing 0.4 M D-sorbitol, 20 mM KCl, 20 mM MES, 10 mM CaCl₂, 0.1% (w/v) BSA, 1.5% (w/v) Cellulase R10 (Duchefa), 0.5% (w/v) Macerozyme R10 (Duchefa), 5% (v/v) Viscozyme® L (Sigma), and 3.56 mM 2-mercaptoethanol. The digestion was carried out for 2 hours at 40 rpm. The resulting protoplast suspension was filtered through a 40 µm mesh and centrifuged at 200 × g for 5 minutes at 4°C using a swinging bucket rotor with minimal deceleration.

Protoplasts were resuspended in 2 mL of washing buffer composed of 0.4 M D-mannitol, 20 mM KCl, 20 mM MES, 1 mM CaCl₂, 0.1% (w/v) BSA, and 3.56 mM 2-mercaptoethanol. Purification of intact protoplasts was achieved using a three-phase density gradient centrifugation with OptiPrep™ (Stemcell Technologies). The OptiPrep working solution (WS) was prepared by dissolving 0.06 g of KCl in 10 mL of OptiPrep. For the gradient, 2 mL of protoplast suspension was mixed with 500 µL of OptiPrep WS (final density: 1.064 g/mL) and layered at the bottom of a 5 mL centrifuge tube. The middle layer consisted of 1 mL of a 2:0.4 (v/v) mix of resuspension buffer and OptiPrep WS (final density: 1.053 g/mL), gently added atop the bottom layer. The top layer comprised 200 µL of resuspension buffer carefully pipetted over the middle layer.

The gradient was centrifuged at 200 × g for 5 minutes at 4°C using a swinging bucket rotor. Protoplasts were recovered from the topmost layer, resuspended in washing buffer, visualized under microscope and adjusted to a final concentration of ∼1,000 cells/μL. The purified protoplast suspension was loaded into the 10x Genomics Chromium Controller for microfluidic droplet generation. Single-cell libraries were prepared using the Chromium Single Cell 3’ Reagent Kit v3.1 (10x Genomics), following the manufacturer’s protocol. Libraries were sequenced on an Illumina NovaSeq X Plus platform (paired-end 150 bp reads).

#### Single cell data analysis

Cellranger (10X Genomics) was used to map the reads to the Arabidopsis reference genome (Araport11). Upon alignment, the ‘force-cells’ option was used to detect the top 10,000 cells based on UMI counts. For each sample, low-quality cells were removed based on mitochondrial and chloroplast read counts (5% mitochondrial and 20% chloroplast counts), which resulted in the identification of 7932 and 5739 high-quality cells in the mock and induced samples, respectively. The Seurat R package (reference) was used for the integration, dimensionality reduction, scaling, clustering and DEG analysis. We used the canonical correlation analysis (CCA) integration option for the integration of both datasets. Uniform manifold approximation and projection (UMAP) were calculated using 32 principal component dimensions and cell-clusters were identified using the Seurat ‘FindClusters’ function with the resolution parameter set to 0.8, which resulted in the identification of 16 cell clusters. For each cluster, gene markers were identified using the ‘FindAllMarkers’ function with a threshold of adjusted P-value < 0.01, log2FC > 0.25, percentage of cells within the cluster that expresses the gene greater than 10% and percentage difference to other clusters greater than 20%. The identities of each cell cluster were determined based on the expression of putative leaf cell-markers ^17^. For the DEG analysis we first aggregate the expression values of both datasets using the ‘AggregateExpression’ function followed by a pair-wise comparison between from mock and treated sample in each individual cluster using the function ‘FindMarkers’, with a threshold of adjusted P-value <= 0.01, log2FC >= 1 for genes that are expressed in at least 20% of the cells.

### Reverse transcription–quantitative PCR (RT–qPCR)

For gene expression analysis, RNA was isolated from protoplast isolated from 4-week-old Arabidopsis leaves 4hpi after E2 infiltration and used for subsequent RT–qPCR analysis. RNA was extracted with a Quick-RNA Plant Kit (R2024; Zymo Research) and treated with RNase-free DNase (4716728001; Merck-Roche). Reverse transcription was carried out using SuperScript IV Reverse Transcriptase (18090050; ThermoFisher Scientific). qPCR was performed using a CFX96 Touch™ Real-Time PCR Detection System. Primers for qPCR analysis of *ISOCHORISMATE SYNTHASE1* (*ICS1*), *ELONGATION FACTOR 1 ALPHA* (*EF1α*) are as follows: *ICS1* primers (*ICS1*_F: 5′-CAATTGGCAGGGAGACTTACG-3’; *ICS1*_R: 5′-GAGCTGATCTGATCCCGACTG-3′). *EF1α* primers (*EF1α*_F: 5′-CAGGCTGATTGTGCTGTTCTTA-3’; *EF1α*_R: 5′GTTGTATCCGACCTTCTTCAGG-3′) and Data were analyzed using the double delta Ct method^40^. All results are plotted using ggplot2 in R^41^, and a detailed statistical summary can be found in Table S1.

### *Albugo candida* propagation and infection assay

For testing the susceptibility in the A. thaliana genotypes in this study, an incompatible isolate of A. candida AcEm2 was used^42^. Infection assays were performed as previously described^32^. Pre-infected leaves with pustules bearing sporangiophores were suspended in water (c. 10^5^ spores ml^−1^). This suspension was incubated on ice for 30 min for release of the zoospores. The inoculum of spore suspension was filtered through a double layered muslin cloth and then sprayed on plants using a Humbrol^®^ (Hornby Hobbies Ltd, Sandwich, UK) spray gun (c. 700 μl per plant). Post spraying, plants were incubated at 4°C overnight in the dark, for promoting zoospore germination. Infected plants were kept under 10 h light (20°C) and 14 h dark (16°C) cycles. Susceptibility on the infected plants with an incompatible strain was scored at 6 days post infection (dpi) by trypan blue imaging of the AcEm2 colonization and its establishment in the host.

#### Trypan blue staining

Infected leaves were boiled for 1 min in stain solution (10 ml lactic acid, 10 ml glycerol, 10 g phenol, 10 mg trypan blue and water in a final volume of 10 ml, mixed in a 1 : 1 ratio with ethanol) and then decolorized in chloral hydrate (2.5 gm chloral hydrate in water in a final volume of 1 ml). The leaves were mounted in 60% glycerol and examined using a Leica M165 FC stereomicroscope.

## Data availability

All sequencing datasets generated in this study have been deposited in the European Nucleotide Archive (ENA) under the accession number PRJEB91113. All custom scripts and analysis code are publicly available at our GitHub repository: https://github.com/dinglab-plants/scETI.

## Author contributions

PD conceptualized and oversaw the inception of the research project. The experimental work was collaboratively conducted by HC, AR, and PD, and. Data analysis and figure generation were performed by LJ, HC, AR, KK, and PD. JDGJ provides valuable discussions that significantly shaped the research during this project. The initial manuscript draft was written by HC, LJ, and PD. All co-authors contributed to subsequent revisions and editorial processes. The final manuscript was prepared by HC and PD and was approved for submission by all authors.

## Acknowledgements

HC and PD acknowledge a European Research Council Starting Grant ‘RELEVATION’ (grant agreement: 101039824). AR acknowledges support by EMBO LTF (ALTF-842-2015). LJ and KK were supported by the Netherlands Organization for Scientific Research (NWO) VIDI grant number VI.Vidi.193.104. JDGJ was supported by the Gatsby Foundation (United Kingdom).

---

**Figure S1.**
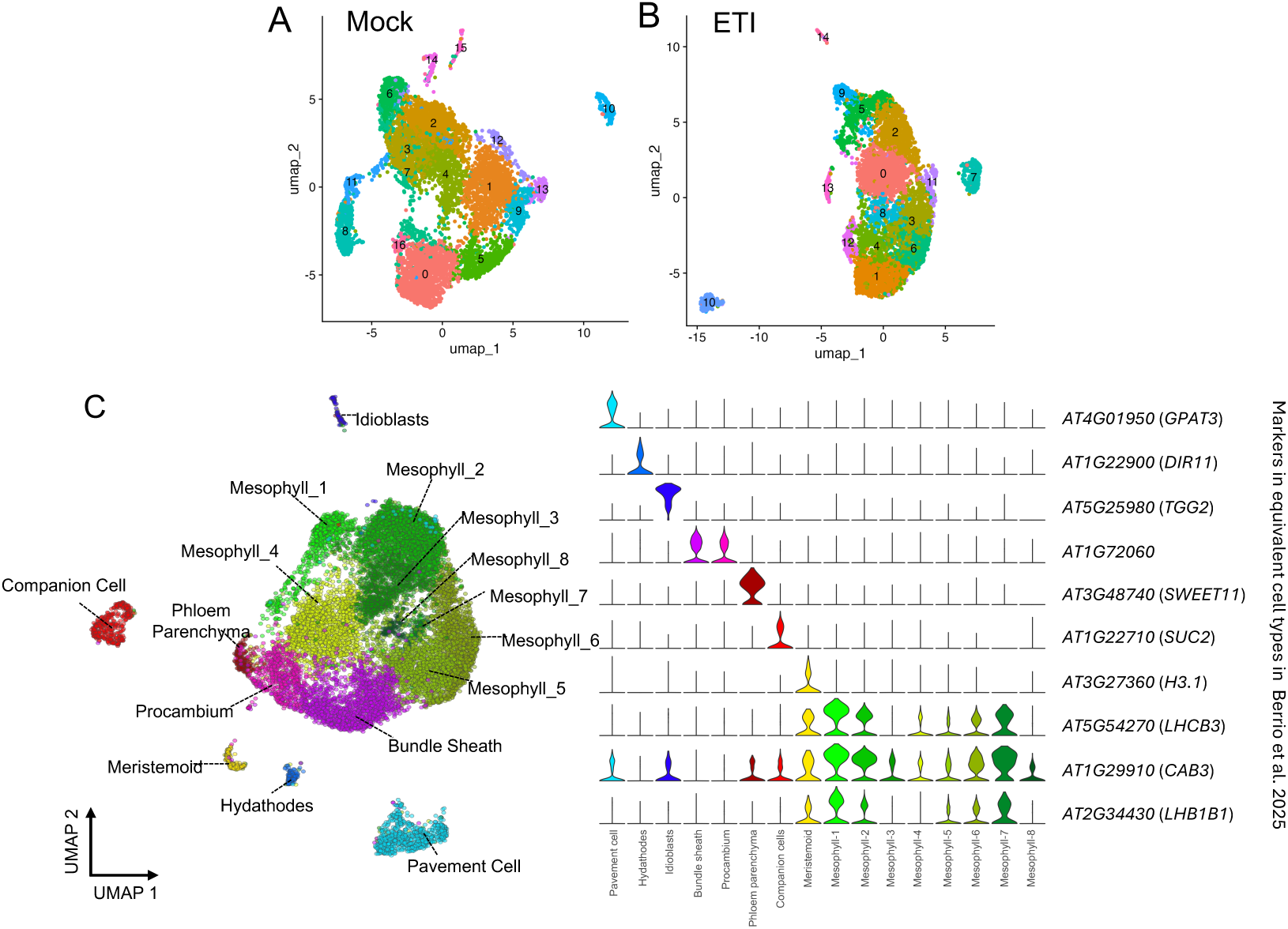
UMAP-based clustering and cell-type annotation of Arabidopsis leaf single-cell transcriptomes. (A, B) UMAP projections of single-cell transcriptomes from mock-treated (A) and ETI-induced (B) Arabidopsis leaf tissues. Cells are colored by unsupervised clusters, corresponding to major and minor leaf cell types. Overall cluster architecture remains conserved upon ETI induction. (C) Cell-type assignment based on the expression of canonical marker genes. Violin plots display representative marker expression across annotated clusters, supporting the identification of pavement, hydathode, idioblast, vascular, mesophyll, and meristem-like cell types. Marker gene identities are cross-referenced with equivalent annotations from Barrio et al., 2025.

**Figure S2.**
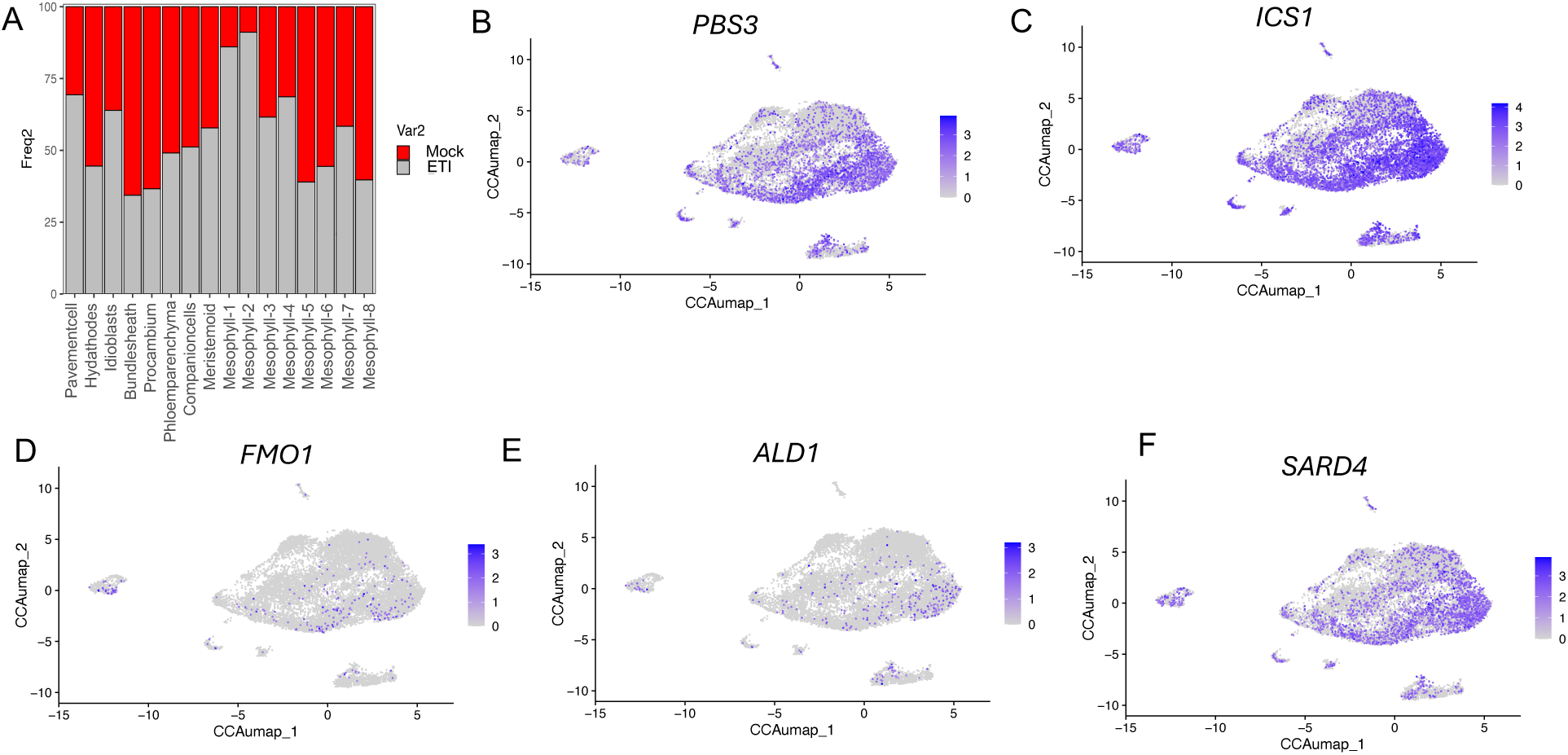
Cell-type compositional changes and activation of salicylic acid and NHP biosynthetic pathways during ETI. (A) Stacked bar plot showing the relative abundance of annotated cell types under mock (gray) and ETI-induced (red) conditions. Increased representation of epidermal and multiple mesophyll subtypes is observed under ETI, indicating shifts in cellular transcriptional states. (B–F) Single-cell feature plots depicting expression of genes involved in salicylic acid (SA) biosynthesis (*ICS1*, *PBS3*) and N-hydroxy pipecolic acid (NHP) biosynthesis (*ALD1*, *SARD4*, *FMO1*). Expression of *ICS1* and *SARD4* is broadly distributed across mesophyll and vascular cells, whereas *FMO1* and *ALD1* show more spatially restricted activation.

**Figure S3.**
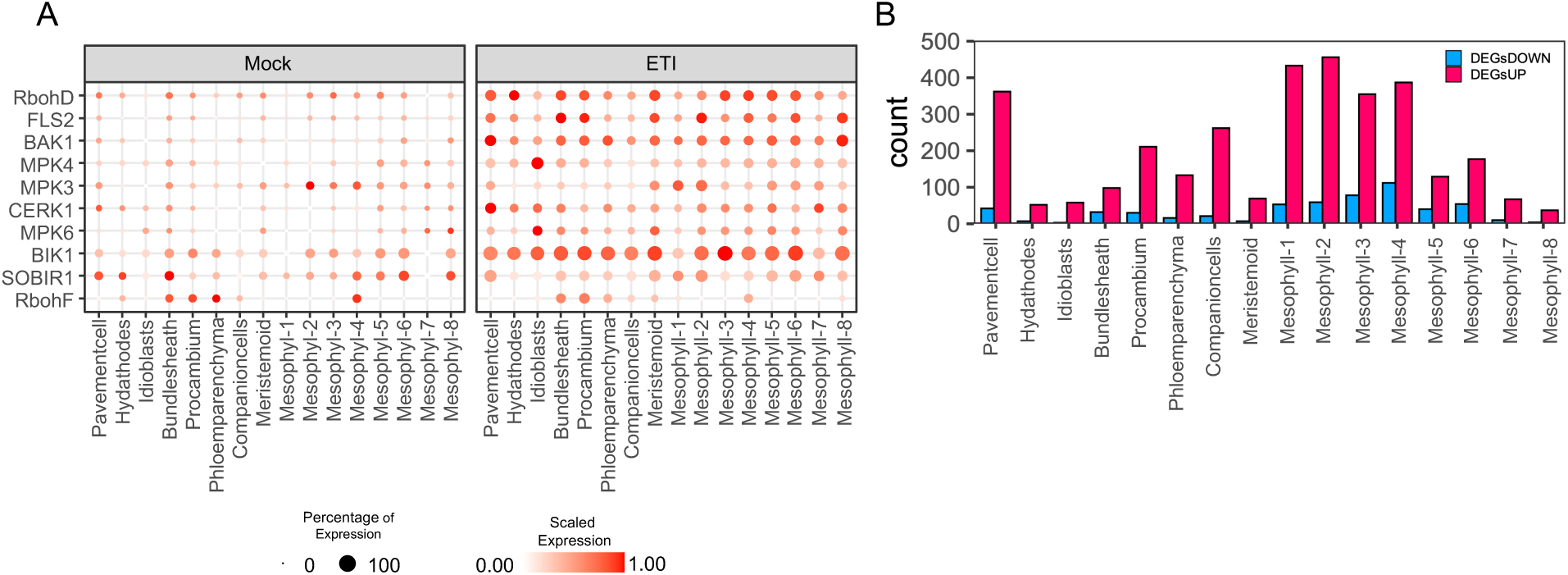
Cell-type-specific expression and transcriptional activation of immune signaling components during ETI. (A) Dot plot illustrating the expression of key pattern-triggered immunity (PTI) and effector-triggered immunity (ETI) signaling genes across annotated leaf cell types under mock and ETI conditions. Dot size indicates the proportion of cells expressing each gene, and color intensity reflects scaled average expression. Genes include receptor kinases (*FLS2*, *BAK1*, *CERK1*), MAP kinases (*MPK3, MPK4, MPK6*), and signaling intermediates (*BIK1*, *SOBIR1*, *RbohD*, *RbohF*). (B) Bar graph showing the number of upregulated (pink) and downregulated (blue) differentially expressed genes (DEGs) across each cell type upon ETI induction. Strong transcriptional activation is observed in mesophyll and pavement cells, consistent with their central role in immune responses.

**Figure S4.**
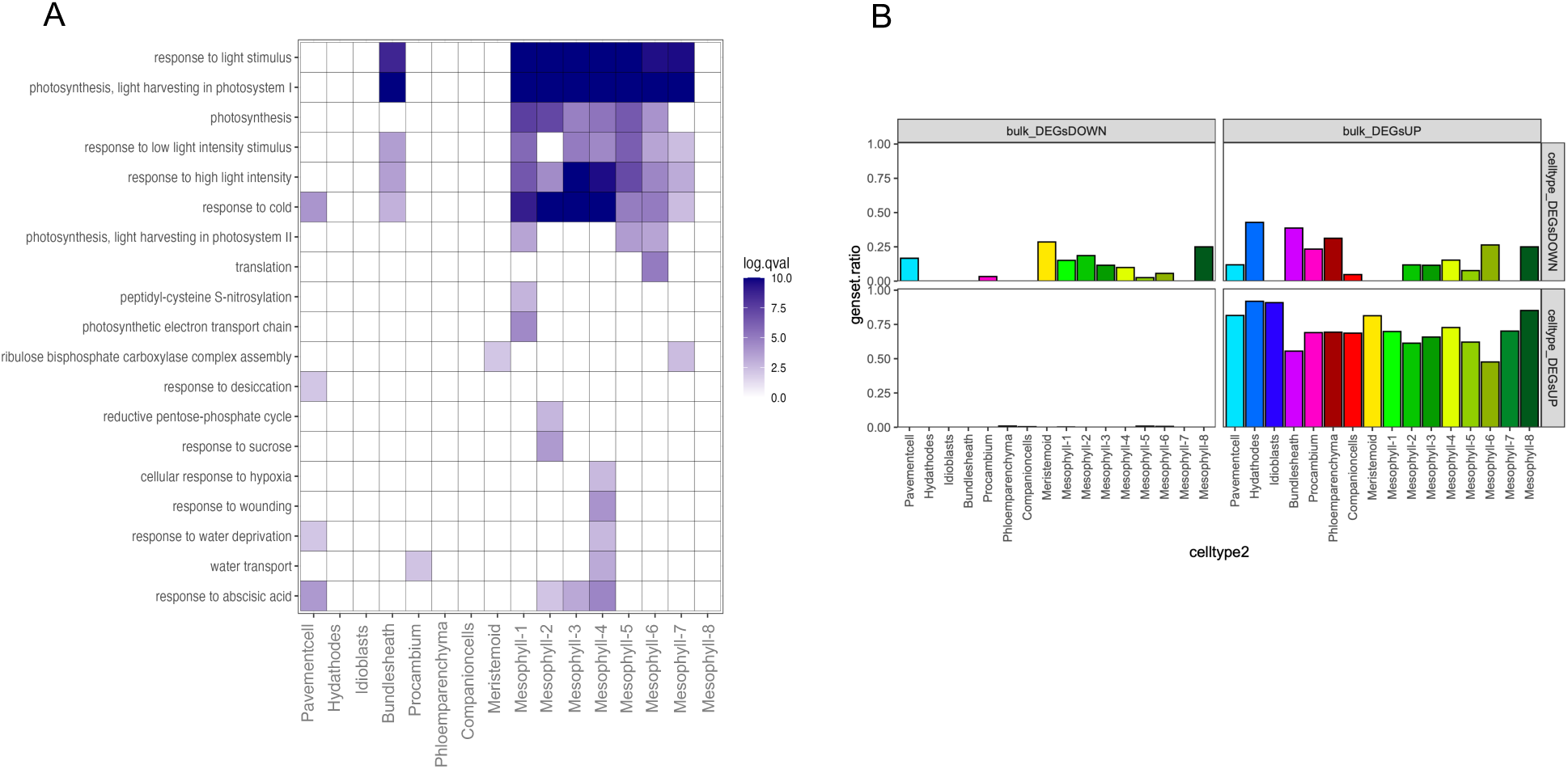
Functional enrichment analysis of bulk ETI-regulated genes across leaf cell types. (B) Heatmap showing Gene Ontology (GO) biological process terms enriched among downregulated genes (DEGsDOWN) mapped to specific cell types using single-cell deconvolution. Color scale indicates statistical significance (log10-transformed adjusted *q*-value). Suppressed photosynthetic and light-responsive pathways are predominantly enriched in mesophyll cluster. (C) Bar plots showing the cell-type distribution of upregulated (DEGsUP, bottom) and downregulated (DEGsDOWN, top) bulk DEGs, inferred by scRNA-seq expression patterns. ETI-upregulated genes are broadly represented across epidermal and mesophyll cells, while downregulated genes are more restricted to photosynthetically active mesophyll populations.

## References

1. Pradeu, T., Thomma, B.P.H.J., Girardin, S.E., and Lemaitre, B. (2024). The conceptual foundations of innate immunity: Taking stock 30 years later. Immunity 57, 613–631. 10.1016/j.immuni.2024.03.007.

2. Sun, T., and Zhang, Y. (2021). Short- and long-distance signaling in plant defense. Plant J. 105, 505–517. 10.1111/tpj.15068.

3. Tabassum, N., and Blilou, I. (2022). Cell-to-Cell Communication During Plant-Pathogen Interaction. Mol. Plant Microbe Interact. 35, 98–108. 10.1094/MPMI-09-21-0221-CR.

4. Jones, J.D.G., Staskawicz, B.J., and Dangl, J.L. (2024). The plant immune system: From discovery to deployment. Cell 187, 2095–2116. 10.1016/j.cell.2024.03.045.

5. Ngou, B.P.M., Jones, J.D.G., and Ding, P. (2022). Plant immune networks. Trends Plant Sci. 27, 255–273. 10.1016/j.tplants.2021.08.012.

6. Zhu, J., Lolle, S., Tang, A., Guel, B., Kvitko, B., Cole, B., and Coaker, G. (2023). Single-cell profiling of Arabidopsis leaves to Pseudomonas syringae infection. Cell Rep. 42, 112676. 10.1016/j.celrep.2023.112676.

7. Tang, B., Feng, L., Hulin, M.T., Ding, P., and Ma, W. (2023). Cell-type-specific responses to fungal infection in plants revealed by single-cell transcriptomics. Cell Host Microbe 31, 1732–1747.e5. 10.1016/j.chom.2023.08.019.

8. Nobori, T., Monell, A., Lee, T.A., Sakata, Y., Shirahama, S., Zhou, J., Nery, J.R., Mine, A., and Ecker, J.R. (2025). A rare PRIMER cell state in plant immunity. Nature 638, 197–205. 10.1038/s41586-024-08383-z.

9. Ngou, B.P.M., Ahn, H.-K., Ding, P., Redkar, A., Brown, H., Ma, Y., Youles, M., Tomlinson, L., and Jones, J.D.G. (2020). Estradiol-inducible AvrRps4 expression reveals distinct properties of TIR-NLR-mediated effector-triggered immunity. J. Exp. Bot. 71, 2186–2197. 10.1093/jxb/erz571.

10. Zhang, Y., Xu, S., Ding, P., Wang, D., Cheng, Y.T., He, J., Gao, M., Xu, F., Li, Y., Zhu, Z., et al. (2010). Control of salicylic acid synthesis and systemic acquired resistance by two members of a plant-specific family of transcription factors. Proc Natl Acad Sci USA 107, 18220–18225. 10.1073/pnas.1005225107.

11. Wang, L., Tsuda, K., Truman, W., Sato, M., Nguyen, L.V., Katagiri, F., and Glazebrook, J. (2011). CBP60g and SARD1 play partially redundant critical roles in salicylic acid signaling. Plant J. 67, 1029–1041. 10.1111/j.1365-313X.2011.04655.x.

12. Sun, T., Zhang, Y., Li, Y., Zhang, Q., Ding, Y., and Zhang, Y. (2015). ChIP-seq reveals broad roles of SARD1 and CBP60g in regulating plant immunity. Nat. Commun. 6, 10159. 10.1038/ncomms10159.

13. Sun, T., Liang, W., Zhang, Y., and Li, X. (2018). Negative regulation of resistance protein-mediated immunity by master transcription factors SARD1 and CBP60g. J. Integr. Plant Biol 60, 1023–1027. 10.1111/jipb.12698.

14. Ngou, B.P.M., Ahn, H.-K., Ding, P., and Jones, J.D.G. (2021). Mutual potentiation of plant immunity by cell-surface and intracellular receptors. Nature 592, 110–115. 10.1038/s41586-021-03315-7.

15. Ding, P., Ngou, B.P.M., Furzer, O.J., Sakai, T., Shrestha, R.K., MacLean, D., and Jones, J.D.G. (2020). High-resolution expression profiling of selected gene sets during plant immune activation. Plant Biotechnol. J. 18, 1610–1619. 10.1111/pbi.13327.

16. Chhillar, H., Nguyen, H.H., Yeh, P.-M., Jones, J.D.G., and Ding, P. (2025). Modular mechanisms of immune priming and growth inhibition mediated by plant effector-triggered immunity. Cell Rep. 44, 115394. 10.1016/j.celrep.2025.115394.

17. Tenorio Berrío, R., Verhelst, E., Eekhout, T., Grones, C., De Veylder, L., De Rybel, B., and Dubois, M. (2025). Dual and spatially resolved drought responses in the Arabidopsis leaf mesophyll revealed by single-cell transcriptomics. New Phytol. 246, 840–858. 10.1111/nph.20446.

18. Duxbury, Z., Wu, C.-H., and Ding, P. (2021). A comparative overview of the intracellular guardians of plants and animals: nlrs in innate immunity and beyond. Annu. Rev. Plant Biol. 72, 155–184. 10.1146/annurev-arplant-080620-104948.

19. Ding, P., and Ding, Y. (2020). Stories of salicylic acid: A plant defense hormone. Trends Plant Sci. 25, 549–565. 10.1016/j.tplants.2020.01.004.

20. Ding, P., Sakai, T., Krishna Shrestha, R., Manosalva Perez, N., Guo, W., Ngou, B.P.M., He, S., Liu, C., Feng, X., Zhang, R., et al. (2021). Chromatin accessibility landscapes activated by cell-surface and intracellular immune receptors. J. Exp. Bot. 72, 7927–7941. 10.1093/jxb/erab373.

21. Ding, P., Rekhter, D., Ding, Y., Feussner, K., Busta, L., Haroth, S., Xu, S., Li, X., Jetter, R., Feussner, I., et al. (2016). Characterization of a pipecolic acid biosynthesis pathway required for systemic acquired resistance. Plant Cell 28, 2603–2615. 10.1105/tpc.16.00486.

22. Hartmann, M., Zeier, T., Bernsdorff, F., Reichel-Deland, V., Kim, D., Hohmann, M., Scholten, N., Schuck, S., Bräutigam, A., Hölzel, T., et al. (2018). Flavin Monooxygenase-Generated N-Hydroxypipecolic Acid Is a Critical Element of Plant Systemic Immunity. Cell 173, 456–469.e16. 10.1016/j.cell.2018.02.049.

23. Chen, Y.-C., Holmes, E.C., Rajniak, J., Kim, J.-G., Tang, S., Fischer, C.R., Mudgett, M.B., and Sattely, E.S. (2018). *N* -hydroxy-pipecolic acid is a mobile metabolite that induces systemic disease resistance in *Arabidopsis*. Proc. Natl. Acad. Sci. USA 115. 10.1073/pnas.1805291115.

24. Hartmann, M., Kim, D., Bernsdorff, F., Ajami-Rashidi, Z., Scholten, N., Schreiber, S., Zeier, T., Schuck, S., Reichel-Deland, V., and Zeier, J. (2017). Biochemical principles and functional aspects of pipecolic acid biosynthesis in plant immunity. Plant Physiol. 174, 124–153. 10.1104/pp.17.00222.

25. Ramirez-Parra, E., López-Matas, M.A., Fründt, C., and Gutierrez, C. (2004). Role of an atypical E2F transcription factor in the control of Arabidopsis cell growth and differentiation. Plant Cell 16, 2350–2363. 10.1105/tpc.104.023978.

26. Wang, S., Gu, Y., Zebell, S.G., Anderson, L.K., Wang, W., Mohan, R., and Dong, X. (2014). A noncanonical role for the CKI-RB-E2F cell-cycle signaling pathway in plant effector-triggered immunity. Cell Host Microbe 16, 787–794. 10.1016/j.chom.2014.10.005.

27. Ayadi, M., Delaporte, V., Li, Y.-F., and Zhou, D.-X. (2004). Analysis of GT-3a identifies a distinct subgroup of trihelix DNA-binding transcription factors in Arabidopsis. FEBS Lett. 562, 147–154. 10.1016/S0014-5793(04)00222-4.

28. O’Malley, R.C., Huang, S.-S.C., Song, L., Lewsey, M.G., Bartlett, A., Nery, J.R., Galli, M., Gallavotti, A., and Ecker, J.R. (2016). Cistrome and epicistrome features shape the regulatory DNA landscape. Cell 165, 1280–1292. 10.1016/j.cell.2016.04.038.

29. Aerts, N., Chhillar, H., Ding, P., and Van Wees, S.C.M. (2022). Transcriptional regulation of plant innate immunity. Essays Biochem. 66, 607–620. 10.1042/EBC20210100.

30. Borhan, M.H., Gunn, N., Cooper, A., Gulden, S., Tör, M., Rimmer, S.R., and Holub, E.B. (2008). WRR4 encodes a TIR-NB-LRR protein that confers broad-spectrum white rust resistance in *Arabidopsis thaliana* to four physiological races of *Albugo candida*. Mol. Plant Microbe Interact. 21, 757–768. 10.1094/MPMI-21-6-0757.

31. Borhan, M.H., Holub, E.B., Kindrachuk, C., Omidi, M., Bozorgmanesh-Frad, G., and Rimmer, S.R. (2010). WRR4, a broad-spectrum TIR-NB-LRR gene from Arabidopsis thaliana that confers white rust resistance in transgenic oilseed Brassica crops. Mol. Plant Pathol. 11, 283–291. 10.1111/j.1364-3703.2009.00599.x.

32. Redkar, A., Cevik, V., Bailey, K., Zhao, H., Kim, D.S., Zou, Z., Furzer, O.J., Fairhead, S., Borhan, M.H., Holub, E.B., et al. (2023). The Arabidopsis WRR4A and WRR4B paralogous NLR proteins both confer recognition of multiple Albugo candida effectors. New Phytol. 237, 532–547. 10.1111/nph.18378.

33. Zimmerli, L., Stein, M., Lipka, V., Schulze-Lefert, P., and Somerville, S. (2004). Host and non-host pathogens elicit different jasmonate/ethylene responses in Arabidopsis. Plant J. 40, 633–646. 10.1111/j.1365-313X.2004.02236.x.

34. Mysore, K.S., and Ryu, C.-M. (2004). Nonhost resistance: how much do we know? Trends Plant Sci. 9, 97–104. 10.1016/j.tplants.2003.12.005.

35. Panstruga, R., and Moscou, M.J. (2020). What is the molecular basis of nonhost resistance? Mol. Plant Microbe Interact. 33, 1253–1264. 10.1094/MPMI-06-20-0161-CR.

36. Cevik, V., Boutrot, F., Apel, W., Robert-Seilaniantz, A., Furzer, O.J., Redkar, A., Castel, B., Kover, P.X., Prince, D.C., Holub, E.B., et al. (2019). Transgressive segregation reveals mechanisms of Arabidopsis immunity to Brassica-infecting races of white rust (Albugo candida). Proc Natl Acad Sci USA 116, 2767–2773. 10.1073/pnas.1812911116.

37. Paauw, M., van Hulten, M., Chatterjee, S., Berg, J.A., Taks, N.W., Giesbers, M., Richard, M.M.S., and van den Burg, H.A. (2023). Hydathode immunity protects the Arabidopsis leaf vasculature against colonization by bacterial pathogens. Curr. Biol. 33, 697–710.e6. 10.1016/j.cub.2023.01.013.

38. Paauw, M., Schravesande, W.E.W., Taks, N.W., Rep, M., Pfeilmeier, S., and van den Burg, H.A. (2025). Evolution of a vascular plant pathogen is associated with the loss of CRISPR-Cas and an increase in genome plasticity and virulence genes. Curr. Biol. 35, 954–969.e5. 10.1016/j.cub.2025.01.003.

39. Wu, F.-H., Shen, S.-C., Lee, L.-Y., Lee, S.-H., Chan, M.-T., and Lin, C.-S. (2009). Tape-Arabidopsis Sandwich - a simpler Arabidopsis protoplast isolation method. Plant Methods 5, 16. 10.1186/1746-4811-5-16.

40. Livak, K.J., and Schmittgen, T.D. (2001). Analysis of relative gene expression data using real-time quantitative PCR and the 2(-Delta Delta C(T)) Method. Methods 25, 402–408. 10.1006/meth.2001.1262.

41. Wickham, H. (2016). ggplot2: Elegant Graphics for Data Analysis (Use R!) 2nd ed. (Springer).

42. McMullan, M., Gardiner, A., Bailey, K., Kemen, E., Ward, B.J., Cevik, V., Robert-Seilaniantz, A., Schultz-Larsen, T., Balmuth, A., Holub, E., et al. (2015). Evidence for suppression of immunity as a driver for genomic introgressions and host range expansion in races of Albugo candida, a generalist parasite. eLife 4. 10.7554/eLife.04550.

